# Cancer-derived Extracellular Vesicles for Targeted Delivery of EGFRvIII siRNA to Glioblastoma, Comparison of siRNA Loading Methods and Efficiency

**DOI:** 10.64898/2026.03.11.710990

**Authors:** Fereshteh Shojaei-Ghahrizjani, Nadim Tawil, Brian Meehan, Laura Montermini, Mohammadrasool Khajeh, Alessandro Maria Villa, Janusz Rak, Paolo Ciana

## Abstract

**Background:** Extracellular vesicles (EVs) are nano and macro-sized, lipid-bound particles, involved in cellular communication. Interestingly, cancer-derived EVs show a heterologous and cross-species tumour tropism which makes them a potential tool for efficient delivery of therapeutic small interfering RNA (siRNA) to the tumour cells.

**Methods:** EVs derived from glioblastoma cells (U373P and U373vIII) were loaded with EGFRvIII siRNA to develop a targeted therapeutic strategy against glioblastoma. EV biodistribution was evaluated using fluorescent indocyanine green (ICG) staining followed by *ex vivo* imaging. Different loading strategies, including passive loading, sonication, saponin-mediated membrane permeabilization, electroporation, and transfection were assessed for their efficiency in loading siRNA into EVs. The efficiency of each method was evaluated by nano flowcytometry, *in vitro* uptake assay followed by immunoblot (western blot) analysis. Eventually, the most effective formulation was tested for the systemic siRNA administration and selective tumour delivery *in vivo*, followed by evaluation of tumour size and EGFRvIII expression.

**Results:** Here, we showed that siRNA transfection into EVs was the most effective loading strategy, as confirmed by nano-flow cytometry, uptake assays, and western blot analysis, achieving over 90% knockdown efficiency *in vitro* for EVs carrying EGFRvIII siRNA. *In vivo*, EGFRvIII siRNA-loaded EVs homed to the tumour site and downregulated EGFRvIII expression compared with the PBS–siRNA control group; however, no significant tumour shrinkage was observed.

**Conclusion:** EGFRvIII-targeting, glioblastoma cell-derived EVs can be used as siRNA delivery carriers for targeted gene therapy in glioblastoma. However, further optimization of siRNA delivery and treatment duration is required.

## 1. Introduction

Glioblastoma multiform (GBM), classified as a grade IV glioma by the World Health Organization (WHO), is the most aggressive and prevalent diffuse glioma of astrocytic origin. Representing 54% of all gliomas and 16% of primary brain tumours, GBM is the most common malignant primary brain tumour (1,2). Despite current treatments, including surgical resection, radiation therapy (RT), and chemotherapy with temozolomide (TMZ), GBM remains incurable, with a median survival of just 15 months (3). Therefore, it is essential to provide more effective treatment approaches, such as precisely targeted delivery of therapeutics, to knockdown specific genes in development of GBM. Epithelial growth factor receptor variant III (EGFRvIII) is the most common mutation in GBM and typically emerges as a late event following EGFR^WT^ amplification (4). EGFRvIII is a smaller receptor due to the deletion of exon 2-7 in the mRNA. As a result, EGFRvIII lacks the ligand-binding domain because of losing amino acids 6–273, making it constitutively active (5). EGFRvIII expression drives cell proliferation, angiogenesis, and invasion across various models and is exclusively found in tumours, not in normal tissues, making it an appealing target for therapies (6–8). EGFRvIII transmits signals through traditional EGFR pathways, such as RAS/MAPK, PI3K/AKT, and JAK/STAT and its tumour-promoting function depends on its kinase activity (9).

In recent years, RNA-based therapies including RNA interference (RNAi) has been widely used. Small interfering RNA (siRNA) is among RNAi techniques which is a post-transcriptional sequence-specific gene silencing method through targeting the desired mRNA (10). Nano-sized vesicles (∼30–150 nm in diameter), with endosomal origin, extracellular vesicles (EVs), have shown significant promise as vectors for delivering therapeutic RNA, such as miRNAs and siRNAs, in cancer therapy (10–12). EVs are highly stable, biocompatible, and often induce low immunogenicity, making them valuable tools for drug delivery and potentially enhancing drug sensitivity and therapeutic outcomes (13).

Some EVs have the tendency to preferentially accumulate in specific tissues, cells, or tumours, which is called EV tropism, often mimicking the properties of their parental cells. This characteristic has garnered significant attention in cancer research, as EVs may serve as natural delivery systems, enhancing targeted treatment strategies. In recent years, the tumour tropism of cancer-derived EVs has been extensively investigated by our group (14–17). By using fluorescently labelled EVs derived from cancer patient plasma (plasma EVs), we demonstrated the potential of autologous EVs for delivering tumour-interfering molecules in a theranostic approach (18). Similar efficacy was observed for EVs derived from large canine breeds with spontaneous brain malignant tumours (19). This feature is not only useful for therapy, but also for intraoperative imaging of tumours through the delivery of diagnostic agents such as indocyanine green (ICG) and Iohexol (17,20).

In this study, we explored various strategies to load glioblastoma cell line-derived EVs with siRNA targeting EGFRvIII. We assessed different exogenous siRNA loading approaches including: (1) passive loading, (2) sonication, (3) saponin-mediated membrane permeabilization, (4) electroporation, and (5) transfection. The efficiency and functionality of each formulation were evaluated experimentally. Finally, a subcutaneous mouse model of glioblastoma was treated with EGFRvIII siRNA-transfected EVs, and their antineoplastic effects were assessed. The results indicate that EGFRvIII siRNA–loaded glioblastoma-derived EVs prepared via transfection can serve as siRNA delivery carriers for targeted gene therapy of glioblastoma.

## 2. Methods

### 2.1 Cells

Human glioblastoma U373 parental cell line (U373P), human glioblastoma U373vIII cell line (containing EGFRvIII mutation: U373vIII) were contributed by the late Dr. Ab Guha (University of Toronto). These cells and human normal astrocyte cell line (NHA) were cultured at humidified incubator with 37°C and 5% CO2 in RPMI-1640 medium (Gibco) supplemented with 10% fetal bovine serum (FBS, Euroclone), 1% of 100 u/mL penicillin/streptomycin (Gibco) and 1% L-glutamine (Gibco). The clonal human epidermoid carcinoma cell line expressing CD63-GFP was prepared by transfecting A431 parental cells with the plasmid pCMV6-AC-GFP (OriGene, Rockville, MD), clone RG 201733 in Dulbecco’s modified eagle medium (DMEM, Gibco) which encodes the fusion protein of CD63-GFP. The brightest clonal sublines (GFP expressing) were then chosen for the experiments (21–23).

### 2.2 EV isolation from conditioned media

To harvest EVs, cells were plated into T-75 flask in 10% FBS containing medium on day 1. On day 2, regular media was exchanged for an EV-depleted 10% FBS containing medium. FBS EV-depletion was carried out using tangential flow filtration (TFF) using 300 KDa molecular weight cut-off hallow fiber filter and KR2i TFF system (Repligen, Massachusetts, USA). The EV-depleted FBS was subsequently filtered with a 0.2 um filter system for sterilization followed by NTA QC to validate efficient depletion of FBS native nanoparticles. Afterwards, EVs were isolated from the conditioned media collected at 72hrs using two different protocols. In method 1, we collected EVs with our standard protocol of production which contains sequential centrifugation steps. It starts with centrifugation of conditioned media at 200 g, 5 minutes followed by 2000 g, 10 minutes and the final ultracentrifugation at 110,000 g for 2h at 4⁰C using 25 ml polypropylene tubes (Beckman, No. 326823). For method 2, after the two centrifugation steps at 200 g and 2000 g, the conditioned media was filtered with 0.8 um syringe filters followed by concentration of conditioned media with 100 KDs molecular weight cut-off Amicon filters (Sigma Aldrich) and finally, an ultracentrifugation at 110,000 g at 4⁰C, using 3 ml propylene tubes (Beckman, No. 349623). Pelleted EVs were washed with PBS and re-pelleted with the same UC settings. We called the method 1 as “non-filtered EVs” and the method 2 as “Filtered EVs”.

### 2.3 Nanoparticle tracking analysis (NTA)

To evaluate the size distribution and concentration of each EV type, NTA was performed using the NanoSight NS300 (Malvern Instruments, Malvern, U.K.). Acquisition was performed with a sample dilution of 1:50-1:100 depending on the sample to achieve a concentration between 20-120 particles/frame. Then, camera level was adjusted at 13, and 5 videos of 60s long were recorded. The videos were then processed using NTA 3.2 analytical software with the threshold level set at 5.

### 2.4 Zeta Potential measurement

Like NTA machine, the ZetaView will record short videos and each particle is counted and tracked in each field of view, giving sample concentration, size and zeta potential. The Zeta potential of EV samples was measured in aqueous buffer using ZetaView PMX120 with ± 200 mV; 1um upper limit of detection.

### 2.5 Immunoblot analysis (Western blotting)

After quantification of proteins with microBCA protein assay kit (Thermo Fisher Scientific, U.S.A.), protein separation was performed using 10% sodium dodecyl sulfate-polyacrylamide gel electrophoresis (SDS-PAGE) (BioRad) followed by transferring to polyvinylidene difluoride (PVDF) membranes. The membranes were then probed with desired primary antibodies, followed by accurate HRP-linked secondary antibodies. Then, the chemiluminescence signal was detected using the Amersham ECL Western Blotting Detection Kit (RPN2108, GE Healthcare) and visualized using the ChemiDoc MP system (Bio-Rad). The antibodies we used in this project were against CD63 (Sigma, catalogue No. SAB4301607, dilution 1:1000), TSG101 (Abcam, catalogue No. AB83, dilution 1:750), ALIX (catalogue No. 186429, dilution 1:1000), CD9 (Abcam, catalogue No. AB223052, dilution 1:750), EGFR (Cell Signaling, rabbit, catalogue No. 4267, dilution 1:1000), EGFRvIII (Cedarlane Labs, mouse, catalogue No. ab00184-1.1, dilution 1:1000), tGFP (Urigene, mouse, catalogue No. TA150041, dilution 1:500), b-Actin (Sigma,catalogue No. A1978-200UL, dilution 1:10,000), anti-rabbit secondary IgG, HRP Linked (Sigma, catalogue No. 7074, dilution 1:5000), anti-mouse secondary IgG, HRP Linked (Cell Signaling, catalogue No. 7076, dilution 1:5000).

### 2.6 ICG staining of EVs

EVs were loaded with Verdye indocyanine green (ICG), as a far-red dye with good *in vivo* penetration with excitation emission 780-814 nm. In summary, EVs were stained by incubating 50 ul EVs (1-5×10^9^ in PBS) with 150 ul of 5 mg/mL ICG in ddH2O, overnight (18h) at 4°C. Next, the samples were transferred to 25 ml polypropylene thin wall tubes (Beckman, NO. 326823), filled up with PBS and ultracentrifuged at 100,000 x g for 92 minutes at 4°C with swing Rotor (SW32Ti, Beckman coulter) to pellet the EVs. The supernatant containing free ICG was removed, and the pellet was resuspended in 50 ul PBS.

### 2.7 Different approaches of loading EGFRvIII Cyanine-5 labelled siRNA into EVs

Our siRNA is specific to the breakpoint caused by the in-frame deletion of exon 2 to 7, which is 21-mer long with the sequence of 5′-CUGGAGGAAAAGAAAGGUAAU-3′ for the sense strand and 3′-GACCUCCUUUUCUUUCCAUUA-5′ for the antisense strand. As mentioned before, for the mere purpose of tracking siRNA, we attached a cyanine 5 at the 3′ of the antisense strand. Cyanine 5 is a red fluorescent dye with excitation of 649 nm and emission of 667 nm. The ability of EGFRvIII siRNA to knock down EGFRvIII expression was first confirmed by a direct transfection of siRNA to U373vIII cells via lipofectamine 2000 transfection reagent. The ratio of siRNA:lipofectamine was optimized as 1:3 and the efficiency of transfection was evaluated after 24h (supplementary figure 1). Second, different methods including passive loading, sonication, saponin-mediated membrane permeabilization, electroporation and transfection were used to load siRNA into EVs as described in the following.

For passive loading of siRNA into EVs, 5×10^9^ EVs were directly mixed with 200 pmol siRNA and kept at 4°C overnight. For permeabilized samples, 5 × 10⁹ EVs were mixed with saponin at a final concentration of 0.125 mg/mL and incubated for 15 minutes at room temperature. The samples were then filtered using 100 kDa Amicon filters to remove saponin, followed by two additional PBS washes. These permeabilized EV samples were subsequently mixed with 200 pMOL siRNA and incubated for 2 hours at 37°C. For sonication, 5×10^9^ EVs were mixed with 200 pMOL siRNA in 150 µL PBS and incubated at room temperature for 30 minutes prior sonication. The samples were then subjected to sonication using a water bath sonicator for 30 seconds, followed by 1 minute of incubation on ice, and a final 30-second sonication step. After sonication, the samples were purified using IZON qEV original size exclusion chromatography columns (qEV single 70 nm, IZON, LTD Science). For electroporation, EVs were mixed with 200 pMOL siRNA in the electroporation buffer (1.15 mM potassium phosphate (pH 7.2), 25 mM potassium chloride). The samples were transferred to a 4 mm cuvette and electroporated at 400 V, 125 μF and ∞ ohms for 10–15 ms using Gene Pulser Xcell Electroporation System (Bio-Rad, Hercules, CA) and then immediately transferred on ice. After electroporation, the samples were purified using IZON qEV original size exclusion chromatography columns (qEV single 70 nm, IZON, LTD Science). Our last method, transfection, was performed according to the EXO-FECT miRNA/siRNA transfection kit instructions (System Bioscience). In summary, 200 pMOL siRNA was mixed with 4 µL of transfection reagent in 80 µL transfection buffer and incubated at room temperature for 15 minutes. Then, 5×10^9^ EVs in 80 µL PBS were added, and the sample was incubated at 37°C for 1 hour. The purification of samples was performed using the provided columns in the kit. Note that for the *in vivo* experiment the siRNA amount for each injection was set to 10 µg of siRNA (equivalent to 800 pMOL) and the same procedure was followed for the scrambled siRNA and PBS-siRNA groups, except no EVs were used for the PBS-siRNA group.

### 2.8 Nano flow cytometry of EVs (CytoFLEX)

To evaluate the fluorescent signal of EVs after testing different approaches to load Cyanine 5-labelled siRNA, the nano flow cytometry (CytoFLEX Flow Cytometer, Beckman Coulter) was used. EVs were diluted in PBS (1:100-1:1000) to have the event rate/sec in the range of 2000-4000 with an error rate below 2%. The proper gating was set after running the unmodified EVs (control EVs) and unmodified PBS, followed by other samples, according to the previously published guidelines (MIFlowCyt, 2008) (24). The fluorescent channel was selected according to the dye. For example, in case of tracking A431 CD63-GFP, the FITC channel was selected and for detection of Cyanine 5, the APC channel was used. The results were then analysed using CytExpert software (version 2.6.0.105).

### 2.9 Uptake assay of EVs

To evaluate the uptake rate of siRNA-loaded EVs via different applied methods, 4×10^5^ U373vIII cells were plated in each well of a 12 multi well plate. Next day, cells were treated with 1.5×10^9^ EVs for 24h. Then, images were captured by fluorescent microscope (EVOS™ M7000 Imaging System, Thermo Fisher Scientific) and flow cytometry analysis was performed (BD FACSVerse) to evaluate the Cyanine 5-positive (siRNA-positive) events indicating the uptake rate of EVs by cells. Flow cytometry analyses of fluorescent cells were performed on 10,000 events on forward scatter (FSC), side scatter (SSC) and the APC channel for detection of Cyanine 5 signal. Results were analysed with BD FACSuite software (BD Biosciences) according to non-treated cells gating to evaluate the EV uptake rate.

### 2.10 Animal model

NSG male, 8-weeks old mice were injected subcutaneously with 2.5×10^6^ U373vIII cells (Animal use protocol number (AUP) at Glen/RIMUHC: 5200). For each experimental group 3 mice were allocated. Mice were monitored daily and eventually, after 17 days, they formed a sizable tumour. The initial tumour size was measured using a standard calliper at day 17 followed by the final measurement at day 25.

### 2.11 Ex vivo imaging of EV biodistribution in a mouse model of glioblastoma

After 24h post injection, *ex vivo* fluorescence imaging was performed using the IVIS Spectrum II Imaging System (PerkinElmer, Waltham, MA, USA) equipped with Cyanine 5.5 filter used for ICG detection. First, mice were anesthetized with 4% isoflurane followed by sacrificing through cervical dislocation and then organs including tumour, liver, lung, brain, heart, spleen and kidneys were collected. Organs were then placed in a 10-cm petri dish with cold PBS followed by imaging over a 6 second exposure time. The quantification of signals was performed via Living Image Software 4.6 (PerkinElmer). For this, the best range of emitted fluorescent signals were set up based on negative controls (mice injected with PBS only) and that range was applied for all images accordingly.

### 2.12 Statistical analysis

All the analysis was performed via GraphPad Prism V.8.2.1 for Windows (GraphPad Software, Boston, Massachusetts, U.S.A). The analytical one-way or two-way ANOVA were performed (based on the dataset) and the significance level was set to p-value <0.05.

## 3. Results

### 3.1 Glioblastoma cell-derived EVs show the tumour tropism

Initially, EVs used in the current study were characterized in accordance with the MISEV2023 international guidelines (25). Specifically, EVs derived from human glioblastoma cell lines (U373P and U373vIII), a non-transformed human astrocyte cell line (NHA), and EVs from a transfected epidermoid carcinoma cell line (A431) expressing CD63-GFP were subjected to comprehensive characterization. As shown in Figure 1A, NTA analysis demonstrated comparable properties among all EV populations, including similar mean and modal particle sizes. The analysis of EV size distribution is also shown in Figure 1B. The expression of Tumour Susceptibility Gene 101 (TSG101) and the tetraspanin CD9 was confirmed by immunoblot (western blot) analysis, while calnexin, used as a negative EV marker, was not detected in EV samples (Figure 1C). In addition, the negative zeta potential of EVs, together with transmission electron microscopy (TEM) analyses, are shown in Figures 1D and 1E, further confirming their EV identity. U373vIII-derived EVs were successfully loaded with indocyanine green (ICG), as assessed by comparison with an ICG standard curve and PBS–ICG controls (Figure 1F), and exhibited an NTA profile comparable to that of unmodified EVs and PBS-ICG control (Figure 1G).

**Figure 1.**
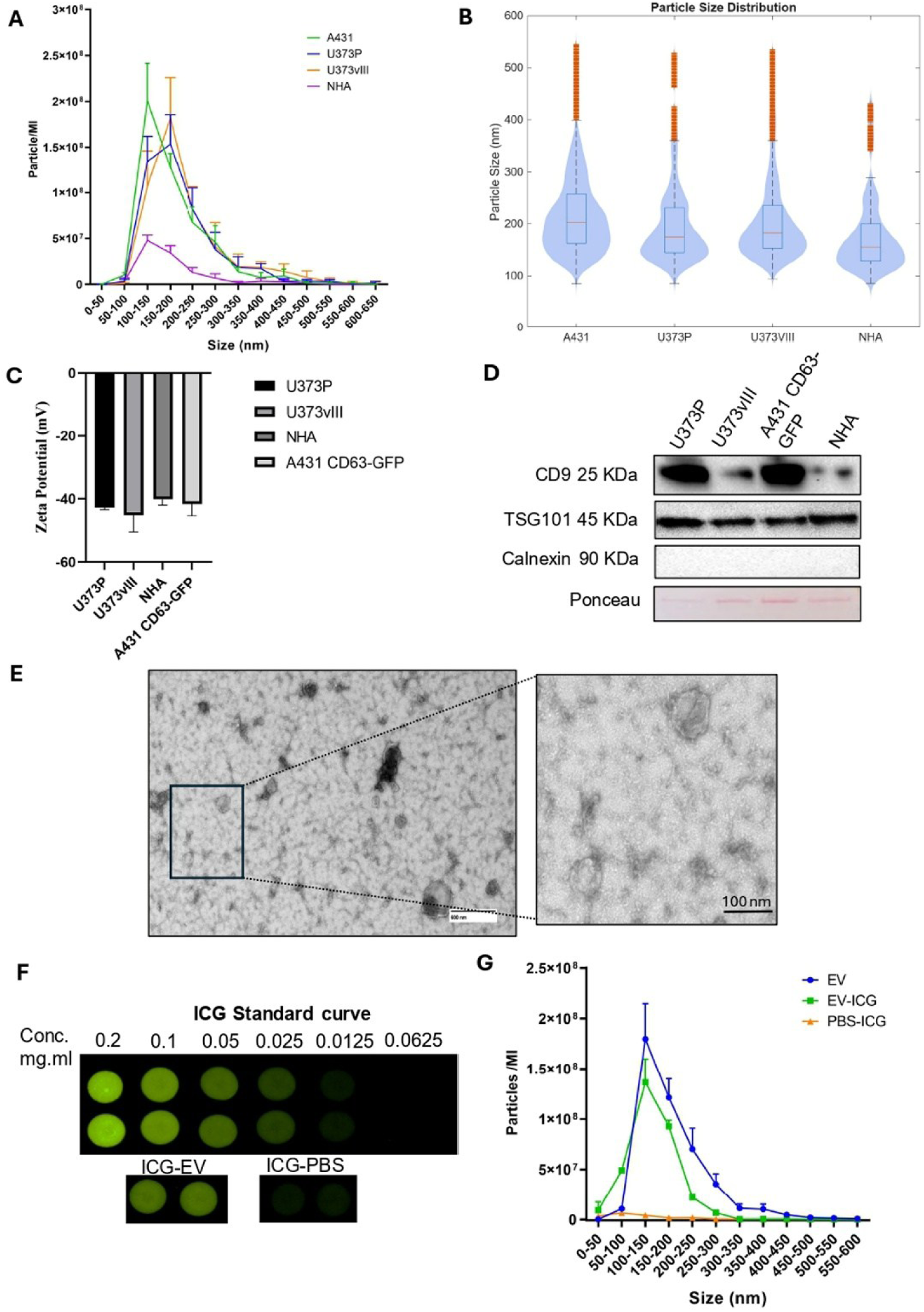
Characterization of extracellular vesicles (EVs). A) Size distribution and concentration of cancer cells-derived EVs including U373 parental, U373vIII, A431 CD63-GFP EVs and normal human astrocyte cells-derived EVs (NHA) shown by NTA. B) Analysis of size distribution profile of EVs, plotted by MATLAB software. C) Immunoblot analysis of EV markers. D) Evaluation of zeta potential of EVs measured by Zeta View. E) Morphology of U373vIII EVs shown by transmission electron microscopy (TEM). F) Quantification of loaded ICG in U373VIII EVs, measured by emitted fluorescent signals of ICG standard curve. G) NTA analysis of ICG-loaded EVs compared to non-modified EVs and ICG-PBS control.

### 3.2 EV isolation protocol defines the *in vivo* biodistribution of EVs

As described in our previous publications, cancer-derived EVs, originating from both cancer cell lines and colorectal cancer patients, exhibit tropism toward neoplastic tissue (15,26–28). We initially compared the tumour tropism of EVs derived from human glioblastoma cell lines (U373P and U373vIII) with those derived from NHA, a non-transformed human astrocyte cell line used as control. A total of 8×10^8^ ICG-loaded EVs were intravenously injected via the tail vein into a subcutaneous mouse model of glioblastoma (n = 3). As controls, mice were injected with PBS–ICG or PBS alone (n = 3). After 24 h, EV biodistribution was evaluated by *ex vivo* imaging using an IVIS Spectrum II instrument. Importantly, differences in EV production using the two methods described in the Methods section resulted in distinct *in vivo* biodistribution profiles. Notably, tumour tropism was observed for ICG-loaded non-filtered EVs (method 1), whereas no accumulation of ICG-loaded filtered EVs (method 2) was detected in mouse tumours (Figure 2A). Quantification of the emitted signals revealed that filtered EVs predominantly accumulated in the liver and lungs, while non-filtered EVs exhibited a higher overall signal intensity. Importantly, U373P-ICG EVs (cancer-derived EVs) showed significantly higher accumulation at the tumour site compared with NHA-ICG EVs and control groups (p < 0.0001) (Figure 2B).

**Figure 2.**
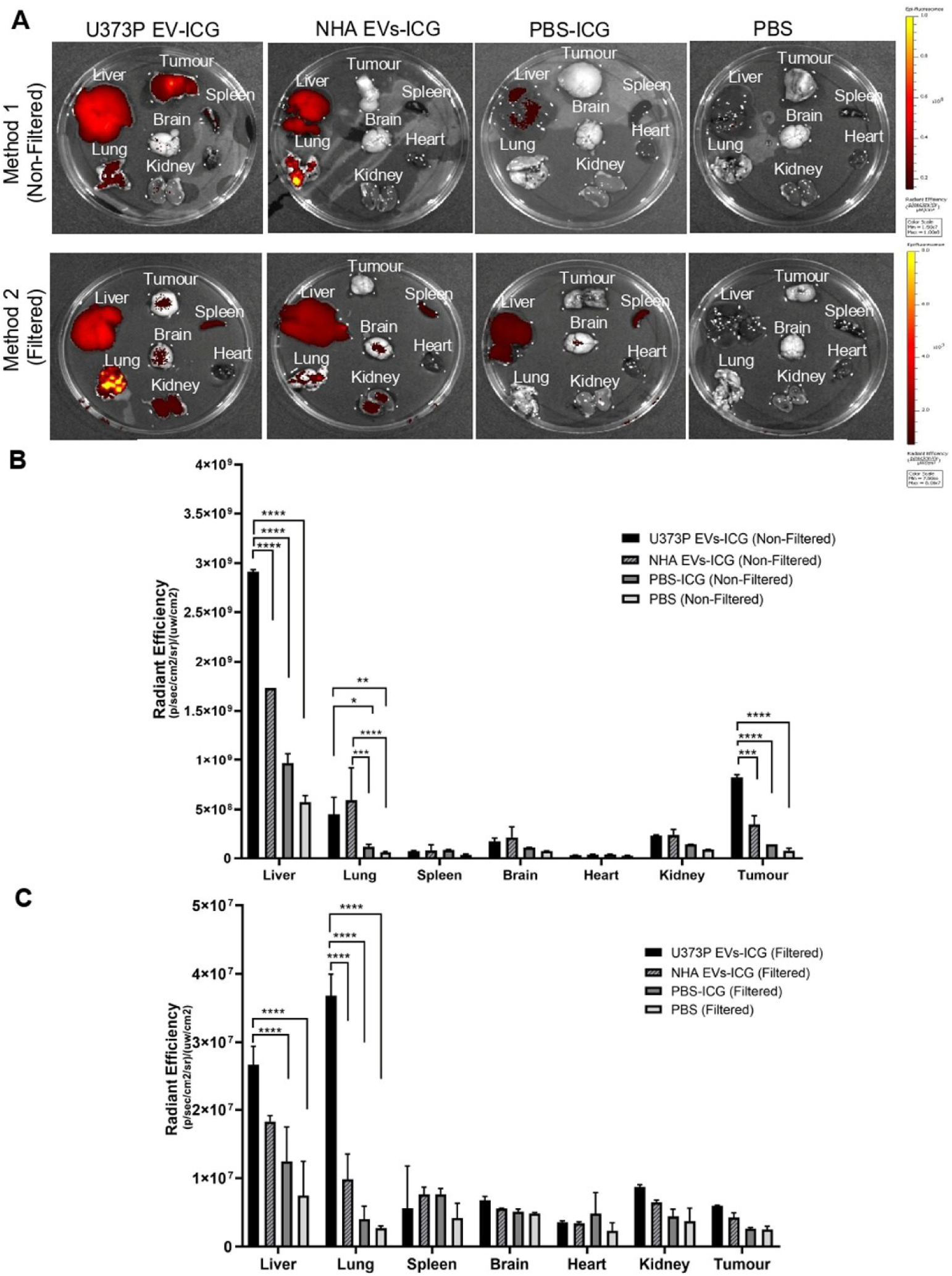
The effect of EV isolation method on distribution profile of cancer-derived EVs in a subcutaneous model of glioblastoma. A) Distribution profile of ICG-loaded U373P EVs, NHA EVs, PBS-ICG and PBS 24h post-injection measured by IVIS Spectrum II, where ICG-loaded EVs were prepared with 2 methods. B) Quantification of the emmitted fluorescent signal from mice 24h post-injection of ICG-loaded EVs prepared via method 1 (Non-Filtered EVs). C) Quantification of the emmitted fluorescent signal from mice organs 24h post-injection of ICG-loaded EVs prepared via method 2 (Filtered EVs). Analysis of experimental groups was performed via two-way ANOVA where significance level has set to p value< 0.05.

### 3.3 Comparison of exogenous siRNA loading strategies for EVs

As mentioned above, exogenous loading methods including passive loading, sonication, saponin-mediated membrane permeabilization, electroporation and transfection were employed to load Cyanine 5-labelled siRNA into A431 CD63-GFP EVs. In nano-flowcytometry analysis, siRNA-loaded EVs were expected to fall within the predefined positive gate in the APC channel. Compared to each corresponding control siRNA-PBS group, only a limited number of positive events was detected for passive loading, sonication, and saponin-mediated permeabilization methods. Although electroporation showed a strong signal in the APC channel, this signal was not considered reliable as a comparable number of positive events was also detected in the corresponding siRNA-PBS control. This has been previously reported in the literature and is attributed to siRNA-induced aggregation (29). In contrast, nano-flow cytometry analysis of EVs loaded via the transfection method revealed a distinct positive population in the APC channel, indicating potential successful siRNA loading compared with the siRNA–PBS control group. It is worth noting that the cyanine-5–positive population shows higher scattering, resulting in a shift of the signal toward larger apparent particle sizes compared with the previously observed profile. To further confirm successful siRNA delivery, an uptake assay was performed. U373vIII cells were treated with siRNA-loaded EVs generated by each method and after 24h, intracellular siRNA levels were evaluated by flow cytometry. Compared with non-treated cells, low levels of positivity were observed for passive loading (0.017%), sonication (0.076%), saponin-mediated membrane permeabilization (2.16%), and electroporation (13.4%). In contrast, EVs loaded via the transfection method achieved the highest uptake efficiency, with approximately 99% positive cells (Figure 3 B, C).

**Figure 3.**
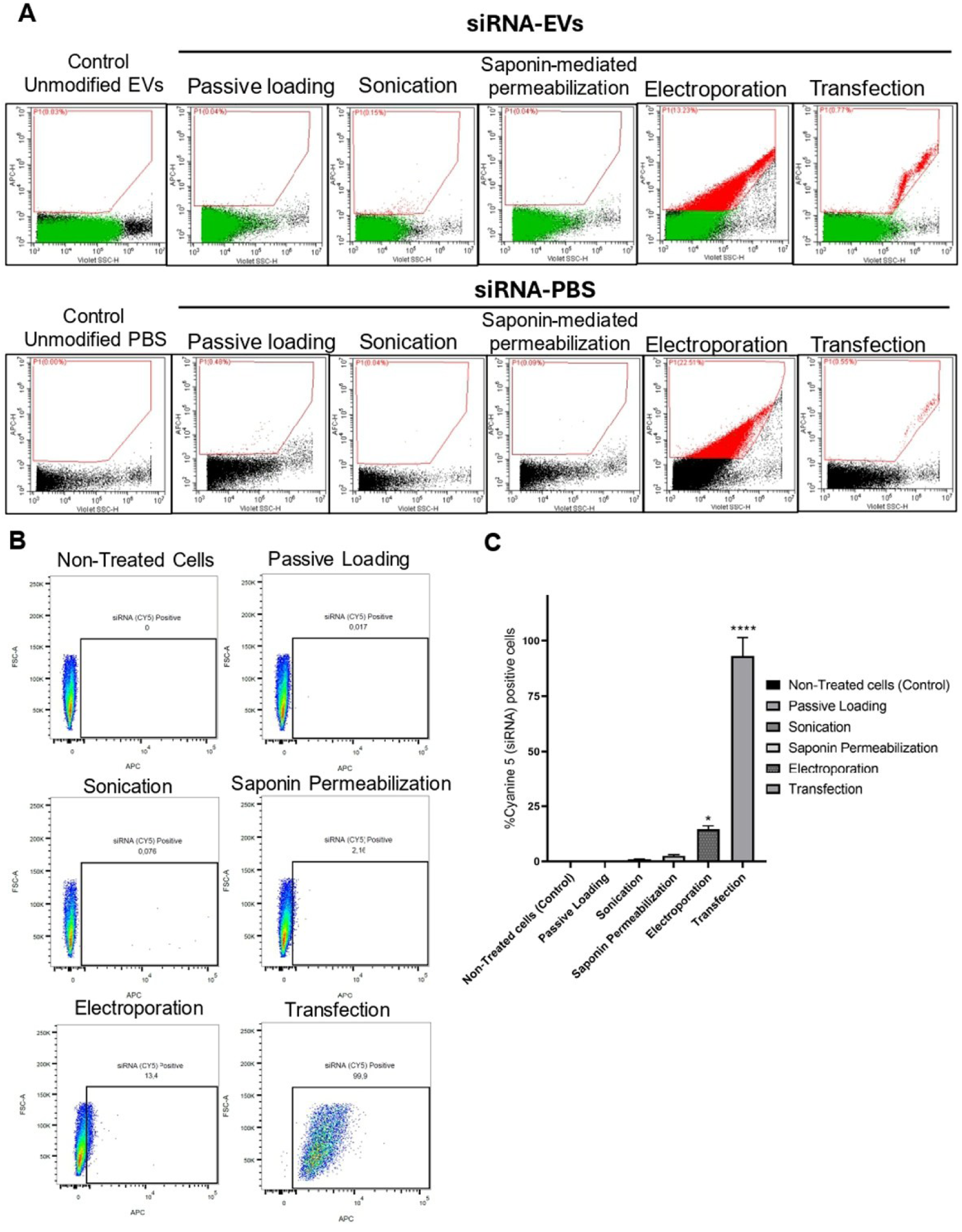
Comparison of siRNA loading methods into EVs. A) Nano-flowcytometry analysis of siRNA-loaded EVs (above) prepared with different methods including passive loading, sonication, saponin-mediated membrane permeabilization, electroporation and transfection compared to siRNA-PBS control (below). The gating strategy was based on control samples of unmodified EVs and unmodified PBS, based on previously published guidelines (24). B) Flowcytometry analysis of treated U373vIII cells with siRNA-loaded EVs (uptake assay) after 24h. The gating strategy was based on non-treated cells’ signal (control) for APC channel. C) Quantification of cyanine 5 (siRNA) positive cells after treatment with siRNA-loaded EVs based on flowcytometry analysis. Analysis of experimental groups was performed via one-way ANOVA where significance level has set to p value< 0.05.

### 3.4 Characterization and functional validation of siRNA-loaded EVs

After selecting transfection as the final method of loading siRNA into EVs, both EV types (A431 CD63-GFP and U373P EVs) were loaded with siRNA. These EVs were characterized by NTA and immunobot analysis for canonical EV markers such as alix, CD9 and TSG101. As shown in figure 4A, the transfection did not alter the EV size distribution profile, and immunoblot analysis confirmed the expression of EV markers in both EV populations (Figure 4B). Interestingly, in the uptake assay, siRNA-loaded A431 CD63-GFP EVs emitted fluorescence signals corresponding to both Cyanine 5-labelled siRNA and GFP, allowing visualization of EV co-localization within treated cells. In contrast, cells treated with siRNA-loaded U373P EVs exhibited fluorescence exclusively from the Cyanine 5-labelled siRNA arising from internalized EVs. The functional efficacy of siRNA-loaded EVs in downregulating EGFRvIII expression was validated by western blot, including appropriate positive and negative controls: cells transfected with siRNA-Lipofectamine and scrambled siRNA, respectively. These results confirmed the ability of siRNA-transfected EVs to selectively reduce the EGFRvIII expression in transfected cells, while the total EGFR expression remained unchanged across all experimental conditions (Figure 4 D, E).

**Figure 4.**
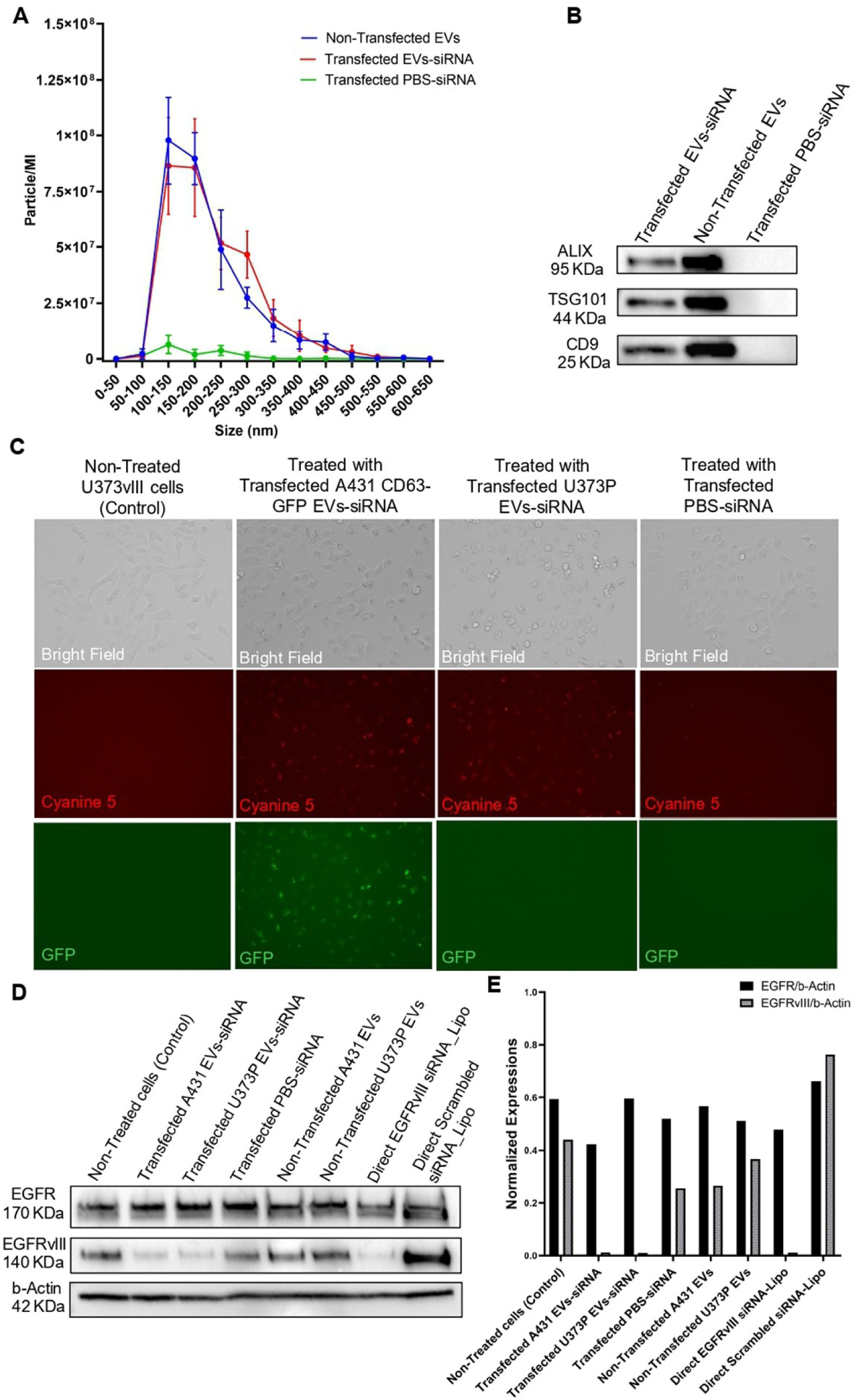
Evaluation of the transfected EVs efficiency in vitro. A) NTA profile of transfected A431 CD63-GFP EVs with EGFRvIII siRNA compared to controls. B) Immunoblot analysis of transfected EVs compared to non-transfected EVs for canonical EV markers. C) Uptake assay of transfected A431 CD63-GFP EVs, U373P EVs and transfected PBS-siRNA by U373VIII cells, 24h after the treatment. Cyanine 5 filed is showing the siRNA positive cells while GFP filed is showing the GFP positive cells. D) Immunoblot analysis of U373VIII cells to evaluate the expression of EGFRvIII and total EGFR 24h after treatment with transfected EVs compared to non-transfected EVs and transfected PBS-siRNA. E) Analysis of EGFR and EGFRvIII to b-Actin expression ratios in different experimental groups measured by Image J.

### 3.5 In Vivo Tumour Targeting and siRNA Delivery by U373vIII-Derived EVs

To evaluate whether siRNA-transfected U373vIII EVs could accumulate in tumours and deliver siRNA to downregulate EGFRvIII, we set-up an *in vivo* experiment (Figure 5 A). Briefly, NSG male mice (8 weeks old) were subcutaneously injected with 2.5×10^6^ U373vIII cells, and after 17 days, palpable tumours of approximately 80 - 150 mm² had developed. As a proof of concept, tumour tropism was first confirmed by comparing the biodistribution of ICG-loaded U373vIII EVs compared to the PBS-ICG control using IVIS spectrum II imaging. The successful accumulation of ICG-loaded U373vIII at the tumour site confirmed their tumour tropism and supported their use for subsequent siRNA delivery experiments (Figure 5 C, D).

**Figure 5.**
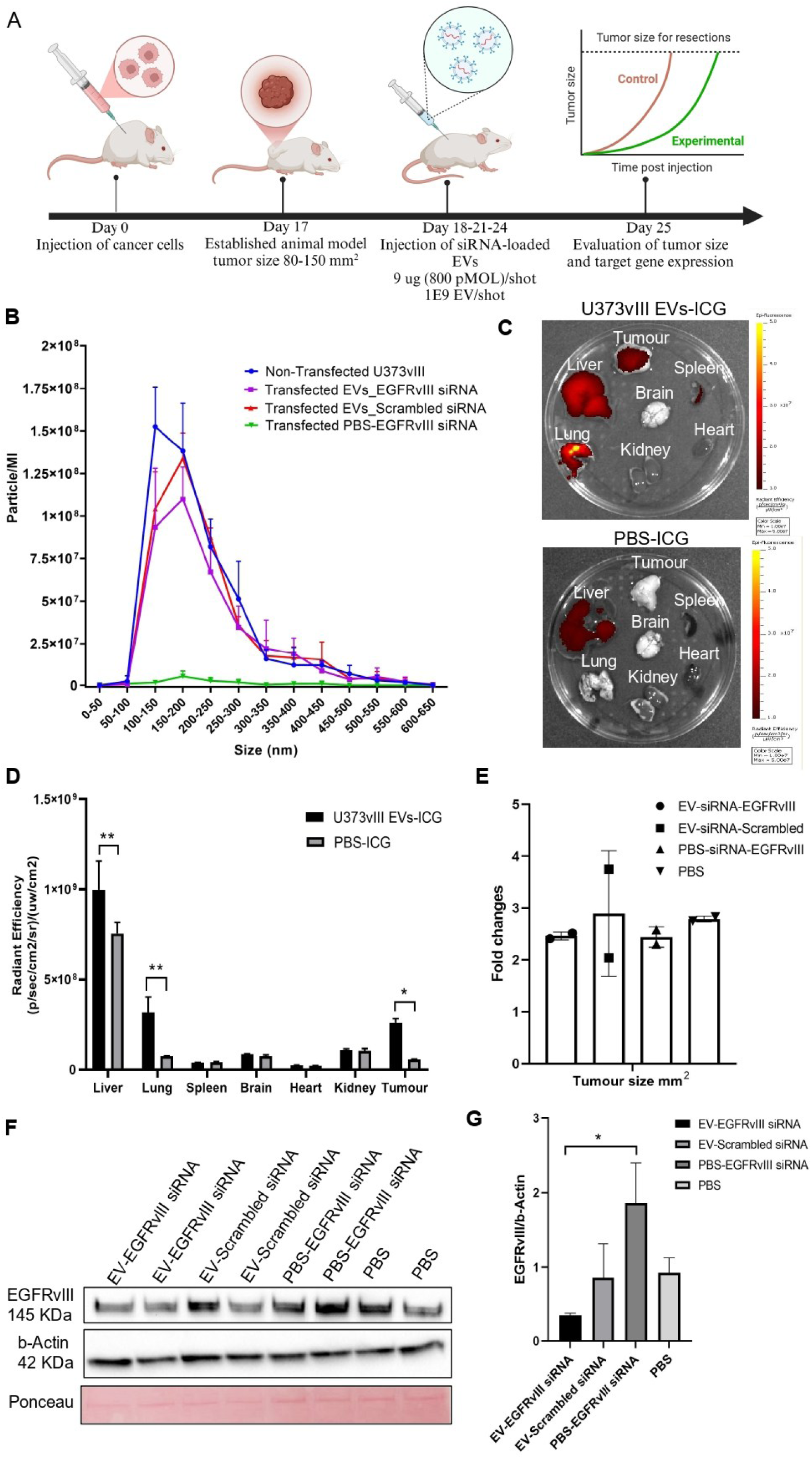
Evaluation of the transfected EVs efficiency in vivo. A) Schematic of the experimental procedure to evaluate the efficacy of transfected U373vIII EVs with EGFRvIII siRNA in a subcutaneous model of glioblastoma compared to transfected EVs with scrambled siRNA, PBS-EGFRvIII siRNA and PBS. B) NTA profile of experimental samples prior injection. C) Biodistribution of ICG-loaded U373vIII EVs *in vivo* compared to PBS-ICG control as a proof of concept. D) Quantification of the emitted fluorescent signal from mice organs and the tumour. Analysis of data was performed via two-way ANOVA, p value<0.05. E) Evaluation of the tumour size for each experimental group, where the ratio of initial tumour size measured on day 17, to the last tumour size measured on day 25, is shown as fold change. F) Immunoblot analysis of tumours to evaluate the expression of EGFRvIII, total EGFR compared to housekeeping b-Actin expression. G) Analysis of EGFRvIII to b-Actin expression ratio in different experimental groups. Analysis of data was carried out with one-way ANOVA, p value<0.05.

To evaluate the *in vivo* efficacy of siRNA-loaded transfected U373vIII EVs, mice were divided into four treatment groups: i) transfected U373vIII EVs with EGFRvIII siRNA, ii) transfected U373vIII EVs with scrambled siRNA, iii) transfected PBS with EGFRvIII siRNA, and iv) PBS alone. Sample characterization by NTA confirmed that the transfection procedure did not alter the EV size distribution profile, and no significant particle formation was observed in the PBS-siRNA control group (Figure 5B). The treatment regimen included three injections administered every 3 days (1×10^9^ EVs/shot for EV groups). Tumour size was measured 24 hours after the third injection, after which tumours were excised for further analysis. Tumour measurements indicated that mice treated with PBS only exhibited the biggest tumour size; however, differences in tumour size among groups did not reach statistical significance (Figure 5 E). In contrast, EGFRvIII expression was significantly reduced in tumours treated with U373vIII EVs loaded with EVs-EGFRvIII siRNA compared to PBS-EGFRvIII siRNA control group (p value<0.05) (Figure 5 F, G). Overall, these results demonstrate the ability of U373vIII-derived EVs to target tumours *in vivo*, and to effectively deliver siRNAs to the tumour tissue.

## 4. Discussion

The intrinsic biological properties of EVs have established them as attractive platforms for the systemic delivery of therapeutic payloads. In the present study, we exploited the innate tumour tropism of glioblastoma-derived EVs to deliver siRNA targeting EGFRvIII, a tumour-specific mutation (6–8). Our findings provide further evidence that cancer-derived EVs can be repurposed as targeted delivery vehicles for siRNA-based therapy, while highlighting the critical impact of isolation and loading methodologies on their functional efficacy.

The observation of preferential tumour accumulation of U373P-derived EVs over non-tumoral astrocyte-derived EVs (Figure 2A) aligns with our previous biodistribution studies (14,15,17,26–28,30–34) which defined this tumour-tropism as heterologous, cross-species, and independent of the tumour type or species from which the EVs originated. While Hoshino et al. (35) and Wu et al. (36) have emphasized the role of integrins and membrane proteins in organ-specific homing, our results reinforce the concept that cancer EVs possess a “self-seeding” capability (37) that can be harnessed for precision medicine. Notably, we observed that the EV isolation process is a determinant of therapeutic success; specifically, the loss of tumour tropism following filtration suggests that such processing may compromise the structural or functional integrity of the EV surface, potentially by stripping essential homing ligands, consistent with earlier reports (38).

A significant challenge in the clinical translation of EV-based therapies remains the identification of an optimal loading strategy. In previous reports, several methods have been proposed, including sonication and electroporation for different types of payloads; in our analysis for siRNA encapsulation, we demonstrated that transfection-based loading was the most effective. Even though electroporation demonstrated high loading efficiency, it generated non-functional aggregates (49), as demonstrated by nano-flow cytometry, that at the end failed to induce functional EGFRvIII downregulation (Supplementary figure 2). Instead, transfection showed a preserved EV marker expression and structural integrity (Figure 4B), thus facilitating successful intracellular delivery and robust gene silencing in recipient cells (Figure 4D). Indeed, in our *in vitro* cell culture and *in vivo* subcutaneous glioblastoma model, administration of siRNA-loaded EVs resulted in significant EGFRvIII downregulation, although we did not observe tumour regression. The lack of a macroscopic therapeutic effect may be attributed to the late initiation of treatment or an insufficient dosing regimen. Nevertheless, the successful molecular targeting of a restricted neoantigen like EGFRvIII *in vivo* represents a proof-of-concept for the use of autologous EV-based delivery. To enhance clinical efficacy, such platforms could be integrated into a multimodal treatment framework, combining them with standard-of-care treatments like temozolomide and radiotherapy, or synergistic gene therapies (e.g., miR-34a) (39). Moreover, the use of EVs as carriers may help overcome the restrictive nature of the blood–tumour barrier formed by glioblastoma-associated vasculature (40) which hinders the efficient penetration of therapeutic agents (41), particularly nucleic acid–based therapies. Our data, together with our previous findings, clearly support the potential of glioma-derived EVs for the targeted delivery of therapeutics across the blood–brain barrier and their accumulation within brain tumours. Collectively, these results highlight EVs as a promising tool to overcome barrier penetration limitations while preserving an intact blood–brain barrier in patients.

In conclusion, our study demonstrates that glioblastoma-derived EVs can be effectively loaded with siRNA to target tumour-specific mutations, including EGFRvIII, showing functional activity *in vitro* and partial therapeutic efficacy *in vivo*. These findings support the feasibility of EV-based siRNA delivery as a personalized RNA interference strategy for EGFRvIII-positive tumours. However, several technical and biological barriers, such as optimization of siRNA-EV dosing, delivery efficiency, tumour microenvironment resistance, heterogeneous signalling pathways, and limitations related to EV isolation, must be addressed to enhance therapeutic outcomes. Future research should focus on refining the therapeutic window, improving delivery strategies, and exploring the synergistic potential of EV-payload combinations in orthotopic models that more closely recapitulate the clinical setting. Overcoming these translational challenges will be critical to fully harness the clinical potential of EV-based siRNA therapies.

## Acknowledgements

This project was supported by grants awarded to P.C. from AIRC (IG 2020 – ID 24914) and from the European Union – NextGenerationEU (PNRR M4C2, Investment 1.4, CN00000041-23, PNRR_CN3RNA_SPOKE8 and SPOKE9), and by grants awarded to J.R. from the Canada First Research Excellence Fund – DNA to RNA (D2R) Research Initiatives, Canadian Institutes for Health Research (CIHR PJT 183971), Fondation Charles Bruneau (FCB) and CIBC, and Canada Foundation for Innovation (CFI10). J.R. is the recipient of the Jack Cole Chair in Pediatric Hematology/Oncology.

We also acknowledge the technical and conceptual support provided by the Centre for Applied Nanomedicine (CAN) at the Research Institute of the McGill University Health Centre (RI-MUHC), Montreal, Canada. https://rimuhc.ca/research-initiatives/centre-for-applied-nanomedicine

## Conflict of interests

The authors declare no conflict of interests.

## Supplementary figures

**Supplementary Figure 1.**
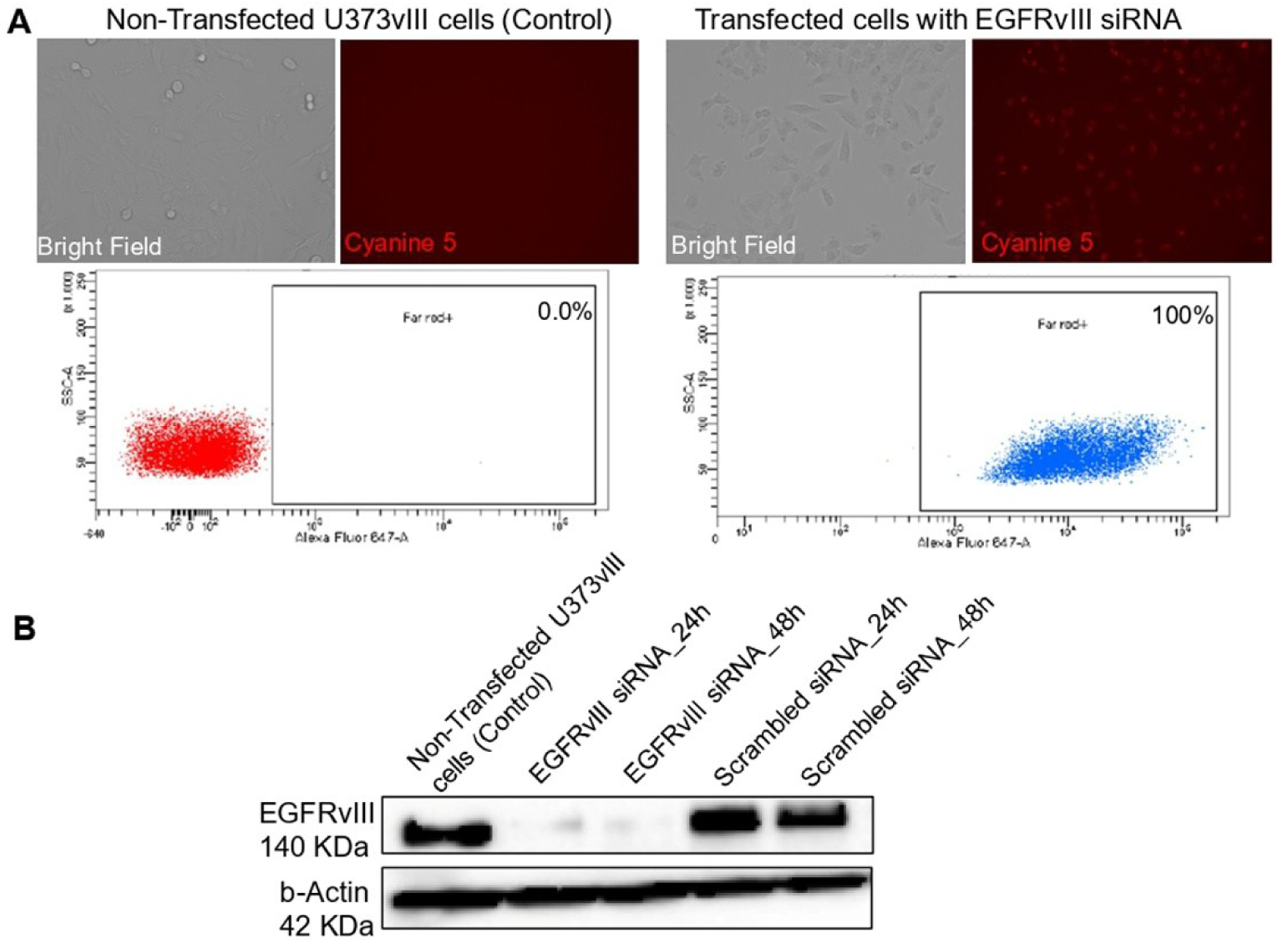
Confirmation of siRNA knockdown efficiency for EGFRvIII expression. A) Imaging of transfected U373vIII cells with siRNA via application of lipofectamine 2000 after 24h compared to non-transfected cells (above). Flowcytometry analysis of siRNA positive cells for non-transfected cells compared to transfected cells (below). B) Immunoblot analysis of EGFRvIII expression in cells 24h and 48h post-transfection with EGFRvIII siRNA, compared to scrambled siRNA.

**Supplementary Figure 2.**
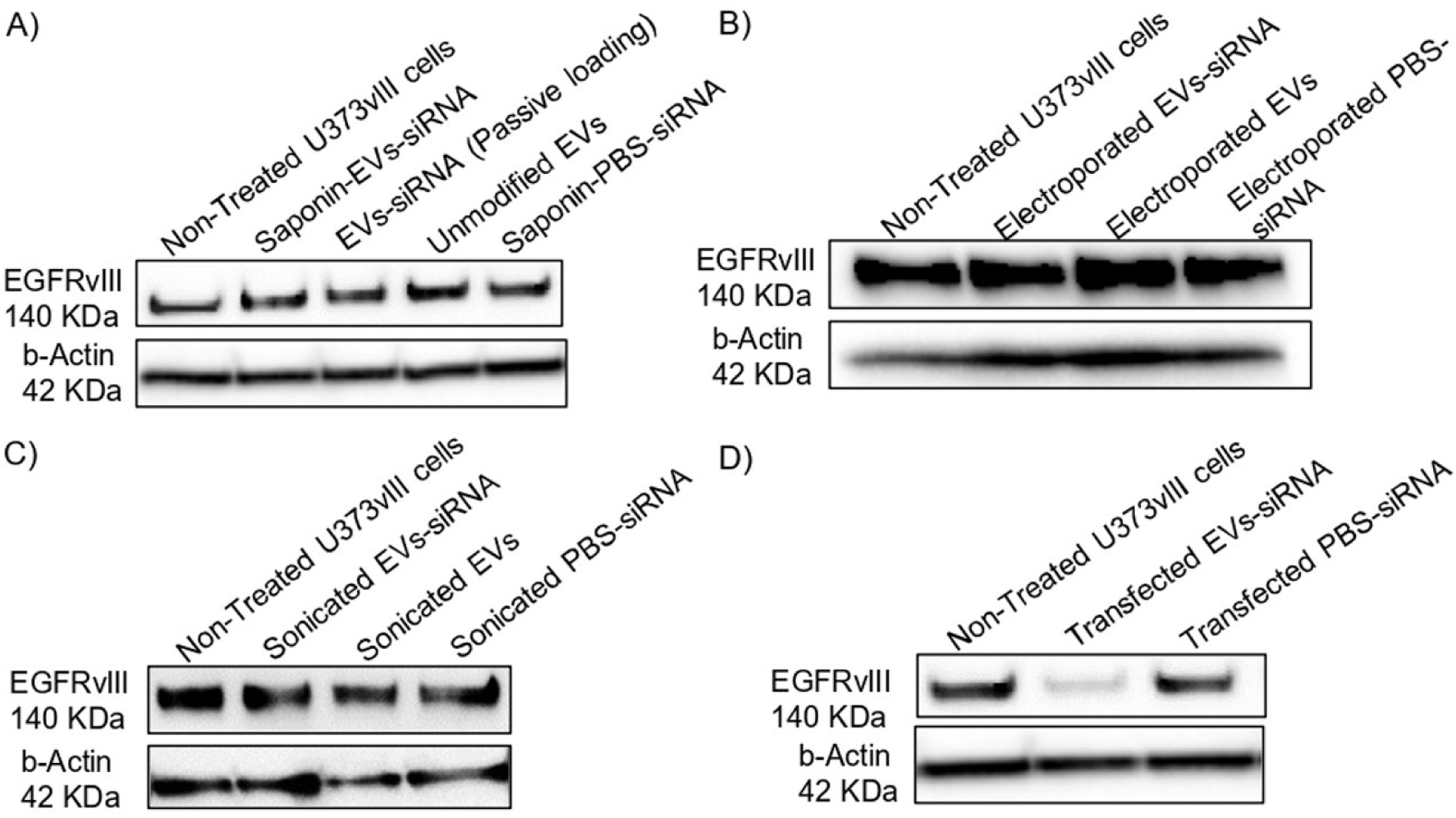
Comparison of EV uptake assay results via immunoblot analysis. U373vIII cells were treated with different formulations and EGFRvIII expression was evaluated 24h after treatment compared to non-treated U373vIII cells (control). A) saponin-mediated permeabilization and passive loading, B) Electroporation, C) Sonication and, D) Transfection.

## Notes

### Competing Interest Statement

The authors have declared no competing interest.

